# Historical trends of the ecotoxicological pesticide risk from the main grain crops in Rolling Pampa (Argentina)

**DOI:** 10.1101/2020.09.02.280560

**Authors:** D.O. Ferraro, F Ghersa, R. de Paula, A.C. Duarte Vera, S. Pessah

**Affiliations:** Universidad de Buenos Aires (UBA), Facultad de Agronomía, Cátedra de Cerealicultura, UBA-CONICET, Instituto de Investigaciones Fisiológicas y Ecológicas Vinculadas a la Agricultura (IFEVA).

## Abstract

We showed the results of the first long-term analysis (1987-2019) of pesticide impact in the main agricultural area of Argentina. Using a clear and meaningful tool, based not only on acute toxicity but also on scaling up the results to total sown area, we identified time trends for both total pesticide impact and the ecoefficiency of modal pesticide profiles. By the end of the time series, soybean showed a pesticide impact four times greater than maize crop in the studied area. However, the time trend in the last years showed a sustainable pattern of pesticide use, with an improvement in the ecoefficiency. Oppositely, maize showed a relatively constant ecoefficiency value during most of the time series, suggesting a possible path towards an unsustainable cropping system. Findings from this study suggest that some efforts have to be made to improve the pest management decisions towards a more efficient pesticide profiles in maize crop and to keep improving the ecotoxicity pesticide profile in soybean crops because of its large sown area in the studied area.

## Introduction

Modern agriculture includes the use of pesticides that have positively impacted cropping systems with a significant increase in yields [1]. However, the potential environmental costs of this intensification process has become a cause for concern [2]. Particularly, rising pesticide use (herbicides, insecticides and fungicides) has been related to both human health and environmental degradation processes [3,4]. Moreover, the global increase of pesticide-resistant organism could lead to a potential rise in pesticide dosage required for the future pest management [5]. Thus, an understanding and a practical assessment of the impact of agrochemical inputs are essential goals for designing sustainable cropping systems [6]. In this sense, a sustainability assessment can be made by using an indicator’s fixed absolute values, or its temporal trajectory, as a proxy for forecasting the future state of a system [7]. The use of long-term approaches has furthered the understanding of the evolution of farming systems [8] and helped to infer future transitions toward sustainable or unsustainable system states [9]. However, data from long-term analyses of pesticide use in the recent literature are scarce [10–12].

There exists an array of indices to measure a pesticide’s toxicity, which provide a hazard assessment of pesticide use with different approaches [13,14]. Almost all these indices are built by combining toxicological data relating to a target pesticide into a single score [11]. However, some indices have several flaws in terms of both transparent comparisons as well as weighting methods [15,16]. A comprehensive assessment requires quantitative indicators as well as system-oriented and diagnostic characteristics. The need to include the above-mentioned aspects in pesticide risk assessment implies the use of quantitative modeling. In addition, models should be able to integrate different types of information, which are not always expressed in the form of empirically based functional relationships but may represent a desirable state regarding the acceptance (or not) of a hazard level.[17].

This paper assessed changes in pesticide risk in the main cropping region of Argentina between 1987 and 2019 using a fuzzy-logic based ecotoxicity hazard indicator [6,18]. In recent decades, the main cropping regions of Argentina have been subject to an intensification process of crop production (i.e. higher yields), as determined by the adoption of no-tillage system [19], the increase in input use (e.g. pesticides and fertilizers), and technological adjustments in crop management (Manuel-Navarrete et al., 2009a; Viglizzo et al., 2003). However, as in the main cropping systems worldwide, the potential impact as well, as the long-term dynamics due to recent technological changes, still remain controversial [20–22]. We analyzed a 32-year period of pesticide use in soybean and maize in the Rolling Pampa (Argentina). In addition, we used crop yields and sown area in order to assess, not only the impact per unit area, but also its possible impact associated with the sown area and the pesticide ecoefficiency (i.e yield achieved per unit of environmental hazard).

## Material and Methods

### Study region

We used data on pesticides, crop yield, and sown area from the Rolling Pampa, the main cropping region of Pampa Region [23]. A pesticide time series was built using the annual profile of pesticides used in the soybean and maize crops. Both crops contributed to 85% and 78 % of total sown area at regional and country level, respectively [24]. The Rolling Pampa is the subregion of the Río de la Plata grasslands with more than 100 years of cropping history [25]. Traditionally a mixed grazing-crop area, the spread of no-tillage in the mid-1990s as well as the wheat–soybean double cropping and the lower cost of inputs (fertilizers, pesticides) led to a rapid expansion and intensification of agricultural production [26]. During the entire long-term period, the changes in the cropping systems of the studied area were mainly represented by three major technological changes: 1) the adoption of no-tillage system (NT); 2) the adoption of genetically-modified organisms (GMO); and 3) the start of systematic fertilization (F). No-tillage minimizes soil mechanical disturbance consequently reducing soil erosion and carbon loss processes, as it leaves a greater percentage of soil covered with plant residues [27]. The change from the conventional tillage system to no-tillage system has also led to a shift in weed control strategy, from a tillage-based scheme to a pesticide-based management strategy. The GMO adoption started in 1996, when the first GMO crop introduced in Argentine agriculture was released, the glyphosate-tolerant soybeans (RR) [28]. The cultivation of RR soybeans, along with transgenic corn hybrids resistant to Lepidoptera (released in 1998) showed an explosive adoption rate among Pampean farmers. It is estimated that 99 % of soybeans and 83% of maize crops in Argentina are GMO [29]

### The fuzzy-logic pesticide indicator (RIPEST)

To assess long-term pesticide hazard dynamics we used RIPEST [6]. RIPEST is a simple fuzzy-based model [30] to estimate the ecotoxicological hazard of pesticides in agricultural systems. The model allows to assess the ecotoxicological hazard for 1) insects, 2) mammals, and 3) the joint hazard of both impacts. The RIPEST structure comprises three main elements: 1) input variables, 2) fuzzy subsets for defining system processes or attributes based on input values and 3) logical nodes for weighting partial indications into a single system performance. The three input variables that describe the toxicity and the amount of active ingredients utilized in each field are: (1) oral acute lethal dose 50 for rats, (2) contact acute lethal dose 50 for bees; and (3) the dose applied for each pesticide application. Therefore, each active ingredient was characterized by means of two different toxicity values: (1) mammal toxicity and (2) insect toxicity. In order to assess the magnitude of the impact of each application, the values of mammal and insect toxicity were measured using the concept hazard quotient [31] defines as:

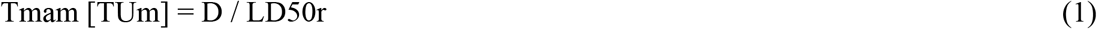

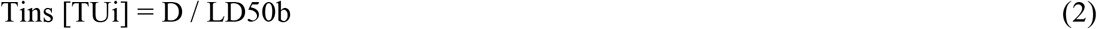

where, Tmam is the mammal toxicity of each pesticide application; Tins the insect toxicity of each pesticide application; D the dose applied (g formulated product/ha); LD50r the oral acute lethal dose 50 for rats (mg formulated product/1000 g rat weight); LD50b the contact acute lethal dose 50 for bees (μg formulated product/bee); and TUi and TUm the toxic units for insects and mammals, respectively. After calculating the Tmam and Tins of single active ingredient formulations and mixtures, RIPEST use the sum of the toxic units (TU) of all the pesticides applied in each field order to calculate the overall toxicity value [32,33]:

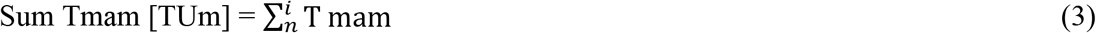

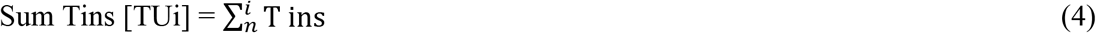

where Sum Tmam is the mammal toxicity of all the pesticides applied; Sum Tins the insect toxicity of all the pesticides applied; and n the number of pesticide applications on each field, during a single cropping cycle. Then, Sum Tmam and Sum Tins values were used to calculate two different indexes: (1) mammal index (M) and (2) insect index (I), according with linear membership in a 0-100 scale. For this scaling, RIPEST uses the Tmam and Tins the highest value for the most toxic pesticides for mammals and insects [6]. These pesticides are Zeta-cypermethrin 0.2 at 200g/ha and Methidathion 0.4 at 1500 g/ha.

Both pesticides involve the highest toxicity registered in the Argentinean National Service for Sanitary and Quality of Agriculture and Food (SENASA 2018) and defines the value of I and M index = 100, respectively. Finally, in order to calculate the overall pesticide impact of pesticides, the (M) and (I) indexes are integrated by two fuzzy rules of the form IF (antecedent)-THEN (consequent) to assemble the pesticide index (P) which indicates the overall impact of pesticide on each analyzed field. P index also range from 0 to 100. In RIPEST the rule node is calculated as follows:

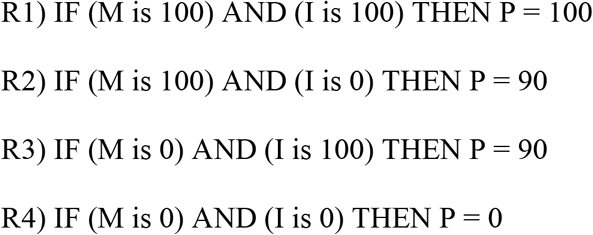

where, R1 to 4 are fuzzy rules; M is the mammal index; I is the insect index; P is the Pesticide index. Finally, the values of all rules are integrated in a single crisp value by defuzzification process using the weighted average method [34].

### Data sources and analysis

Data on pesticide use per hectare were defined following prescriptions for each crop during the whole studied period based on a national reference publication “Márgenes Agropecuarios” (http://www.margenes.com) for all years from 1987 to 2019. Crop yields and sown area were extracted from the Agricultural Estimates of the Ministry of Production and Labor of the Argentine Republic (http://datosestimaciones.magyp.gob.ar/) for the same time period. As the tillage system has shifted from conventional to no-tillage regime, different pesticide profiles were registered for each tillage systems in each crop in the time interval 2002-2007 (soybean) and 2002-2012 (maize). During this period, total values were calculated using data of sown area under these two different tillage systems [35,36]. Before and after these periods each crop under different tillage systems share the same pesticide profile. Final P index value is expressed in a 0-100 scale and represents the ecotoxicological hazard of pesticides applied per hectare. In order to scale the P values for the total sown area, we represent the P index using units (units. ha −1). The pesticide ecoefficiency (i.e yield achieved per unit of environmental hazard) was calculated as the cost–benefit ratio of the yield to environmental impacts. To detect possible monotonic trends in the time series we used the Mann-Kendall test [37]. Pesticide data used in the analysis have been provided as supplementary information.

## Results

Pesticide index (P) showed both different values and time trends in soybean and maize crops (Fig. 1). Soybean showed the highest P value at the beginning of the studied period and decreased until the early-2000, while maize crop started the time series showing extremely low P values (Fig. 1).

**Fig 1.**
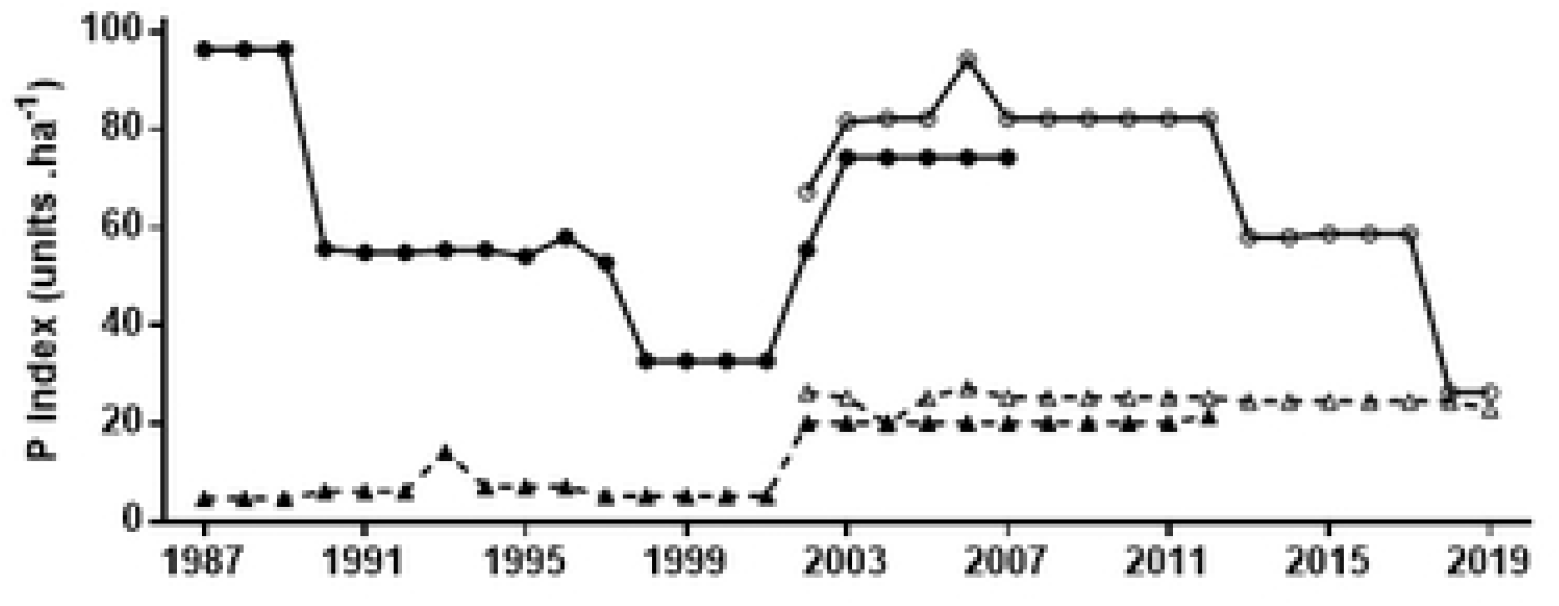
P index [P units. ha −1] for soybean (solid line) and maize (broken line) cropping systems from 1987 to 2019. Time series also shows temporal trend of P index for cropping systems under conventional tillage (CT: closed symbols) and no-tillage (NT: open symbols) for both crops. Mann-Kendall test for monotonic trend: Soybean CT (tau = −0.06, P = 0.69); Soybean NT (tau = −0.33, P = 0.06). Maize CT (tau = 0.63, P = 0.002); Maize NT (tau = −0.53, P = 0.006).

However, by the end of the period, the soybean crop showed a remarkable P index decrease, resulting in similar values for both crops in the last year analyzed (Fig. 1). The high ecotoxicological hazard per hectare (i.e P index) that the soybean crop exhibited during most of the time series was enhanced by the large sown area occupied by this crop in the studied area (Fig. 2).

**Fig. 2.**
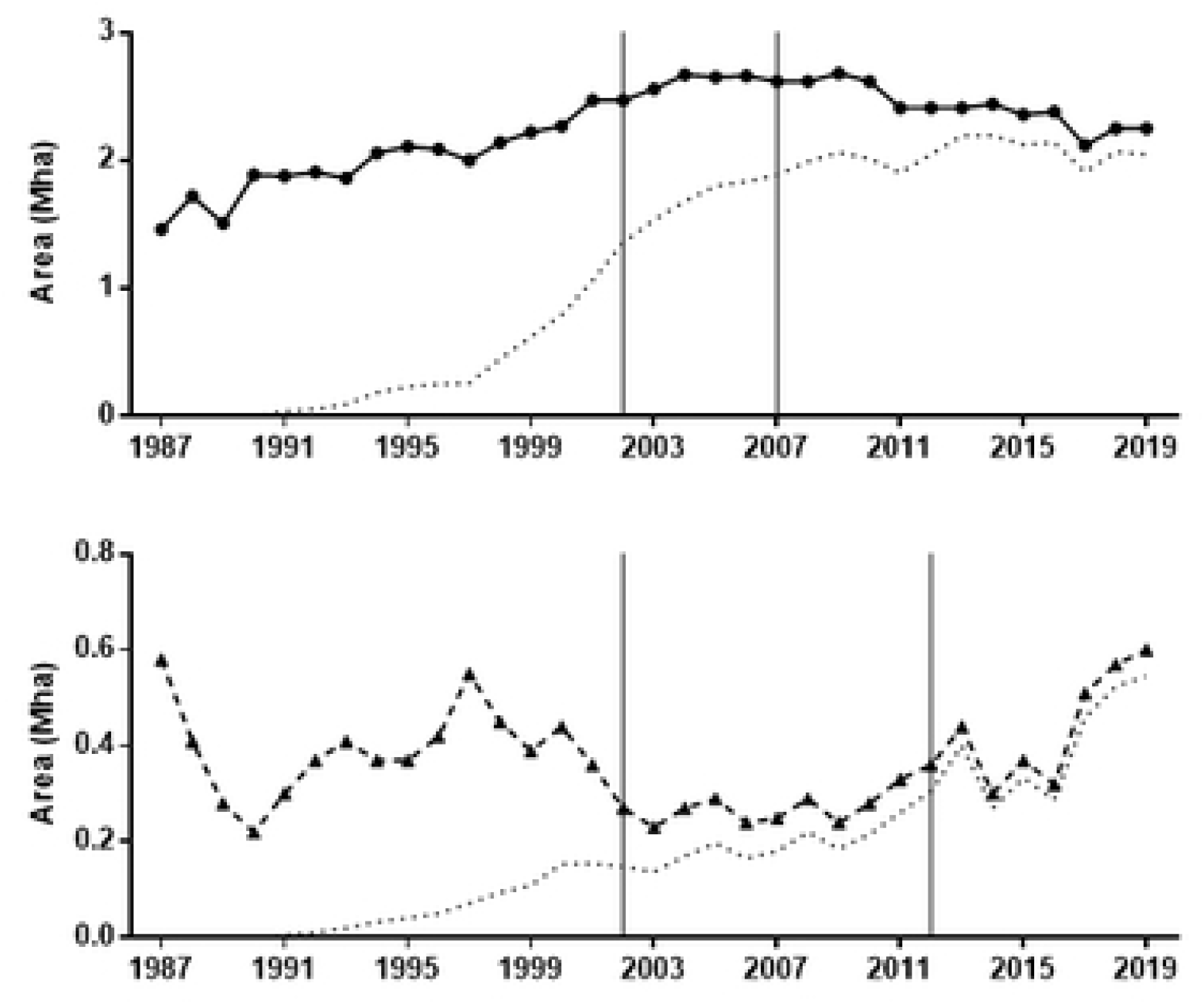
Total sown area (Mha) under soybean (upper panel) and maize (lower panel) crops from 1987 to 2019. The dotted lines show the evolution of area under no-tillage during the studied period. The vertical lines define the time period when the sown area of each crop exhibited differential pesticide usage according to the tillage system (CT and NT). Mann-Kendall test for monotonic trend of total area: Soybean (tau = 0.48, P < 0.001); Maize NT (tau = −0.02, P =0.79).

The increase in the sown area with soybean crop was significant in the period 1987-2019, something that did not occur with corn, which exhibited increases and decreases without a defined pattern (Fig. 2). When P index values were scaled by sown area of each crop, soybean values were one order of magnitude higher than maize in most of the period studied (Fig. 3).

**Fig. 3.**
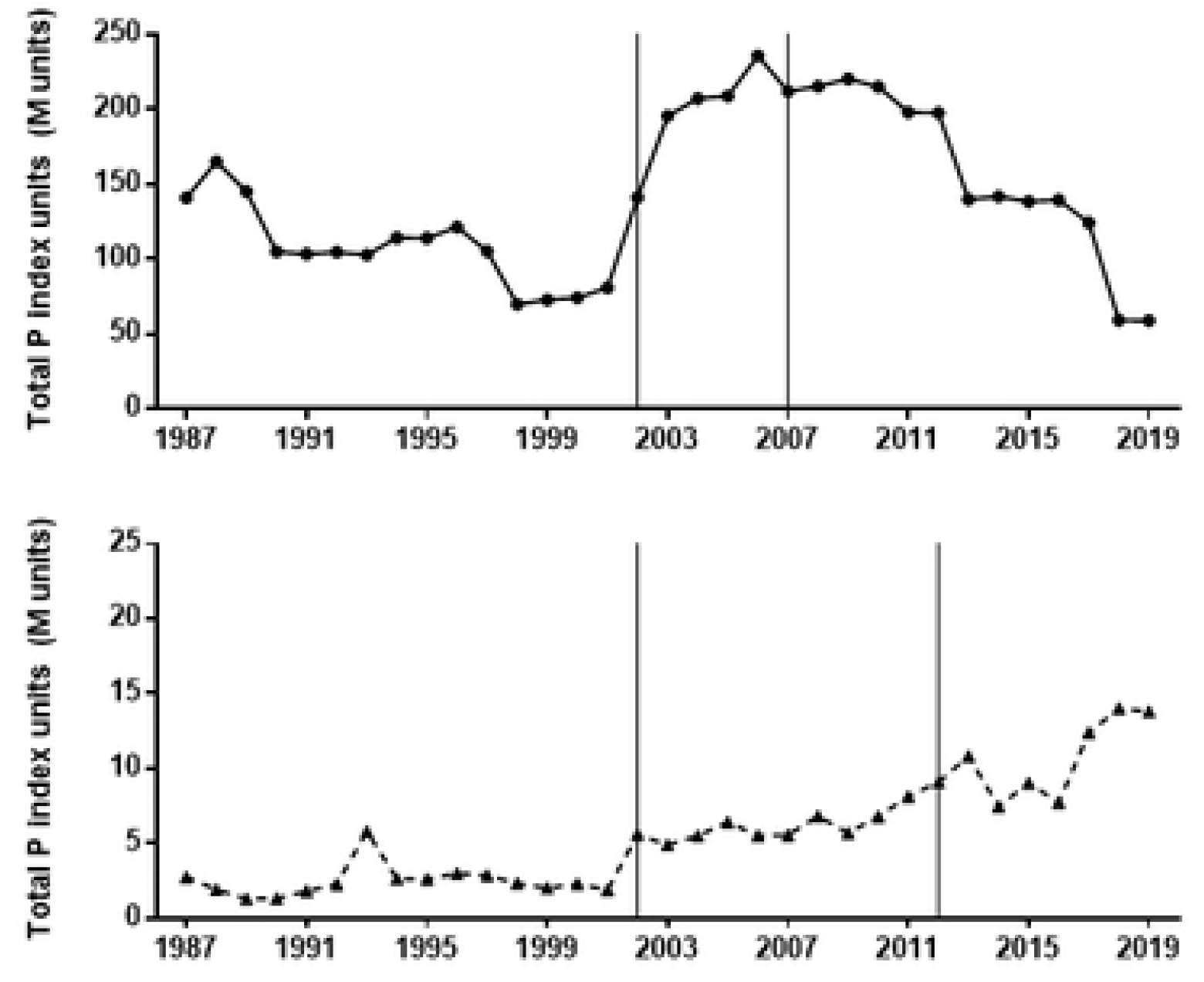
Total P index units [Million of units] under soybean (upper panel) and maize (lower panel) crops. Mann-Kendall test for monotonic trend: Soybean (tau = 0.16, P = 0.20); Maize (tau = 0.71, P < 0.001). The vertical lines define the time period when the sown area of each crop exhibited differential pesticide usage according to the tillage system (CT and NT).

There were also trend differences between crops as soybean showed no significant increase in total P units and maize exhibited a significant increase of P units from 1987 to 2019. The decreasing trend observed in soybean P index (Fig. 1) explained the gap narrowing between the impacts of soybean and maize, which still remained wide by the end of the period, as soybean and maize showed 59 M and 13 M units, respectively (Fig. 3). Both crops showed a monotonic increasing trend in total yield in the studied area (Fig. 4).

**Fig. 4.**
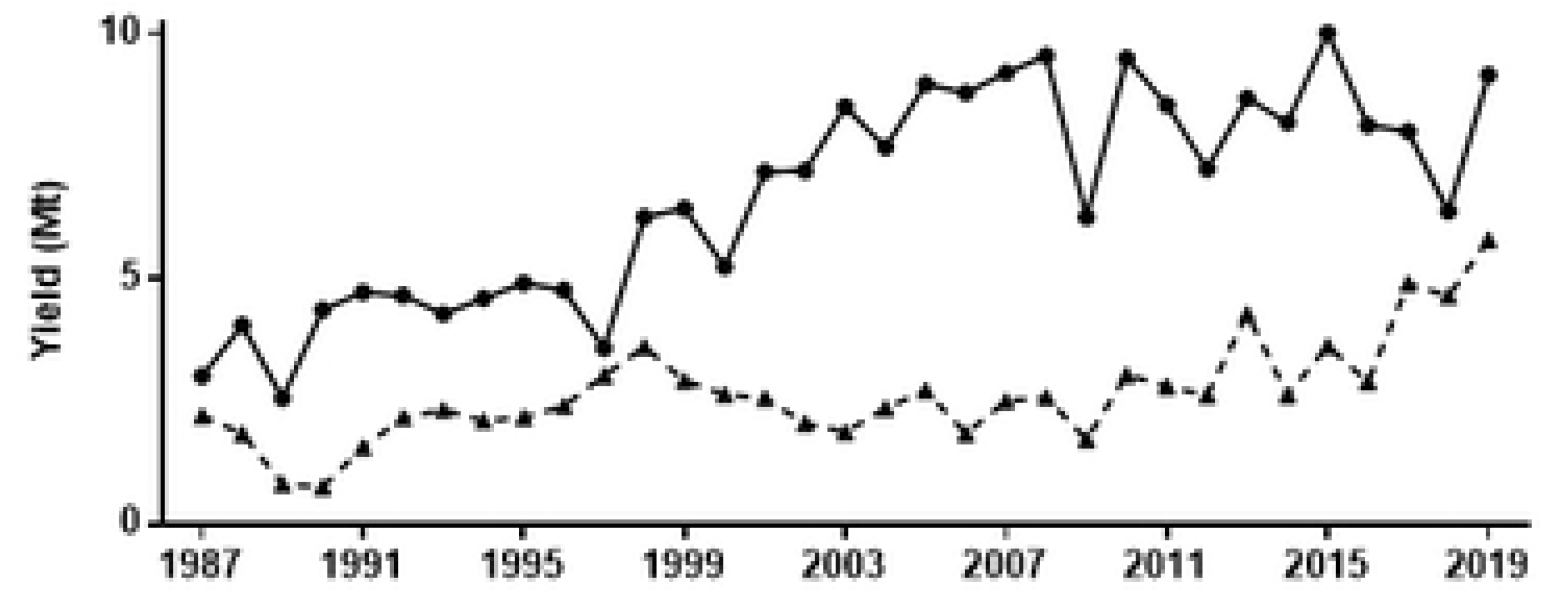
Total yield (Mt) under soybean (solid line) and maize (broken line) crops from 1987 to 2019. Mann-Kendall test for monotonic trend: Soybean (tau = 0.61, P < 0.001); Maize (tau = 0.47, P < 0.001).

However, when both total impact (Fig. 3) and yield (Fig. 4) were integrated in a single ecoefficiency indicator the time trends were different between crops (Fig. 5). Soybean crop showed a significant positive trend in ecoefficiency, with a remarkable improvement from the early-2010 (Fig. 5). Oppositely, maize crop showed an overall negative trend in ecoefficiency, mainly related to a significant decrease when the pesticide profile of no-tillage began to be considered (Fig. 5).

**Fig. 5.**
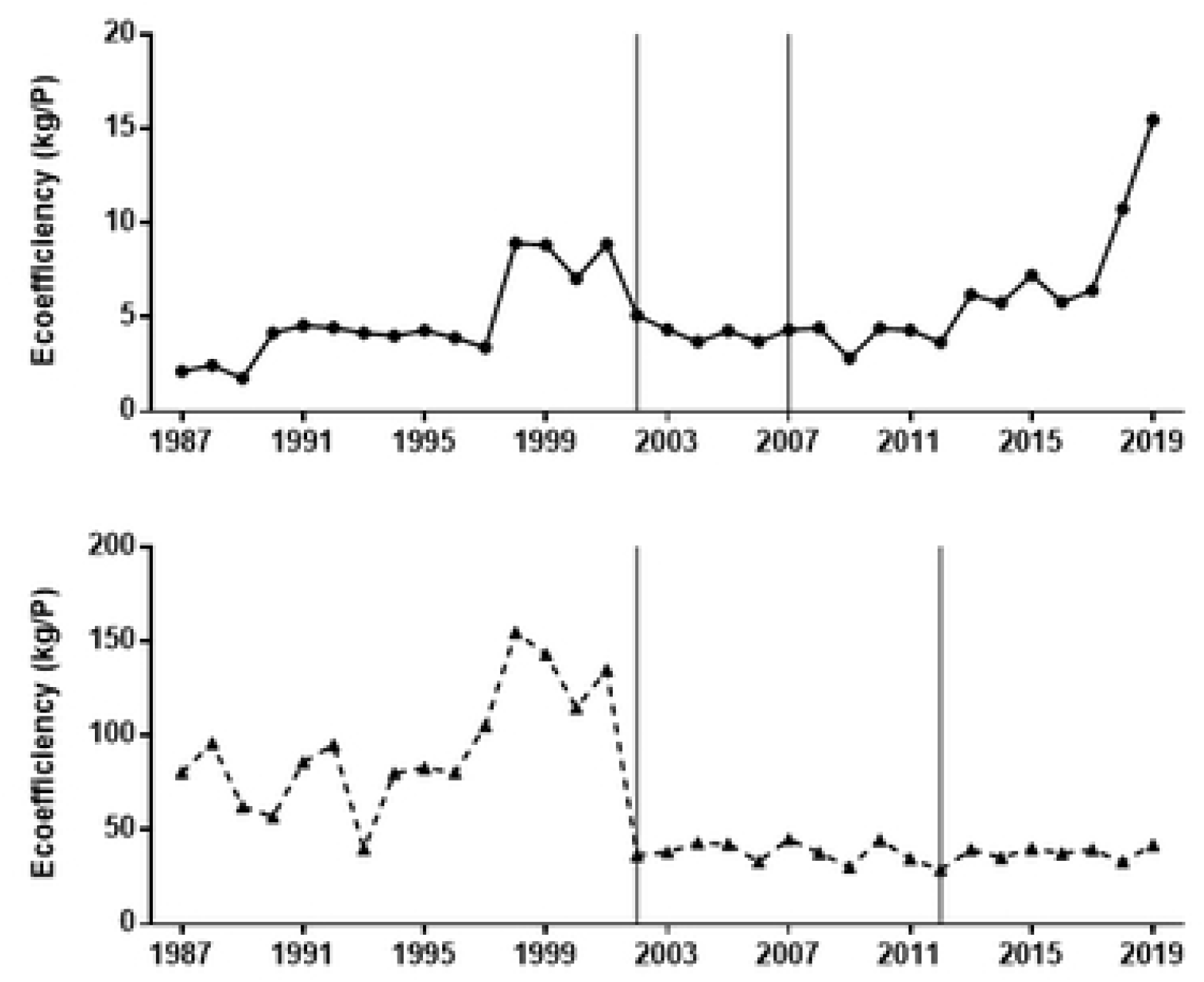
Ecoefficiency (t/P) of soybean (upper panel) and maize (lower panel) crops from 1987 to 2019. Ecoefficiency is based upon the ratio of crop yield to P index. Mann-Kendall test for monotonic trend: Soybean (tau = 0.31, P = 0.004); Maize (tau = −0.44, P < 0.001). The vertical lines define the time period when the sown area of each crop exhibited differential pesticide usage according to the tillage system (CT and NT).

In a relative time-trend analysis of both total pesticide impact (Fig. 4) and the ecoefficiency (Fig. 5) using a common base (1987=1), soybean crop showed no significant trend during the studied period for either measure, but maize showed an almost six-fold increase in total impact, measured as the total number of P units (Fig. 6).

**Fig. 6.**
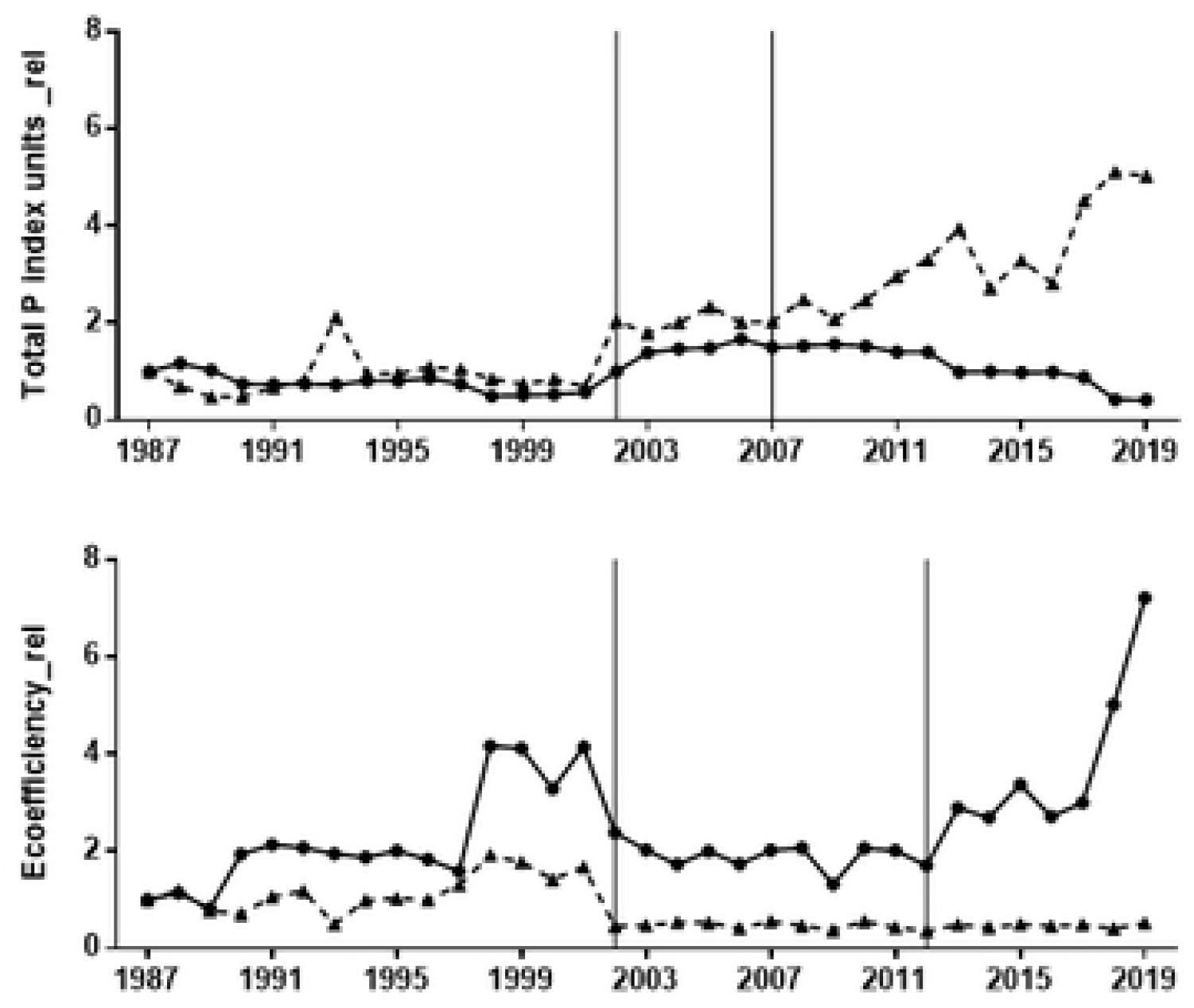
Relative values (Base 1987 =1) of Total P index units (Total P index units_rel: upper panel) and Ecoefficiency (Ecoefficiency_rel: lower panel) of soybean (solid lines) and maize (broken lines) crops from 1987 to 2019. Mann-Kendall test for monotonic trend: Soybean Total P index units_rel (tau = 0.16, P = 0.20); Maize Total P index units_rel (tau = 0.70, P < 0.001); Soybean Ecoefficiency_rel (tau = 0.31, P = 0.004); Maize Ecoefficiency_rel (tau = −0.44, P < 0.001). The vertical lines define the time period when the sown area of each crop exhibited differential pesticide usage according to the tillage system (CT and NT).

When ecoefficiency was analyzed in relative terms, soybean and maize crops not only showed opposite time trends (Fig. 5) but also different magnitudes in these changes. As maize showed a negative trend, in relative terms, reaching a relative value of 0.52 by the end of the period, ecoefficiency of the soybean crop remarkably increased, showing a relative final value of 7.22 in 2019 (Fig. 6). Pesticide profiles in both crops showed noticeable changes in the 1987-2019 period in the quantity of active ingredient used as well as in the specific toxicity of the pesticides (Tables 1 and 2).

**Table 1.**
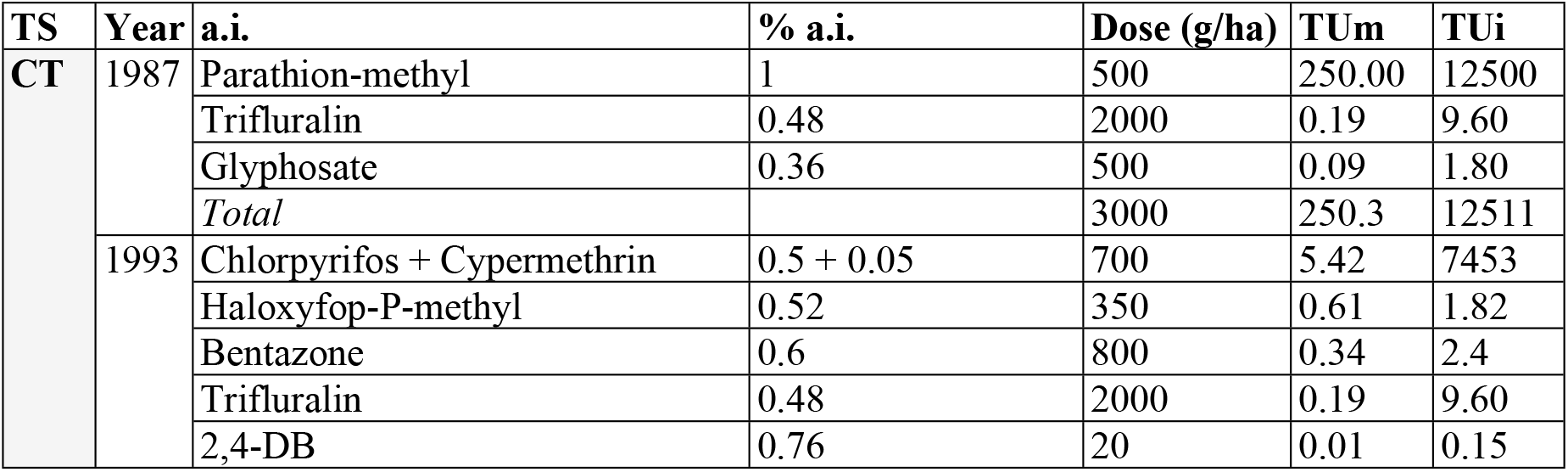
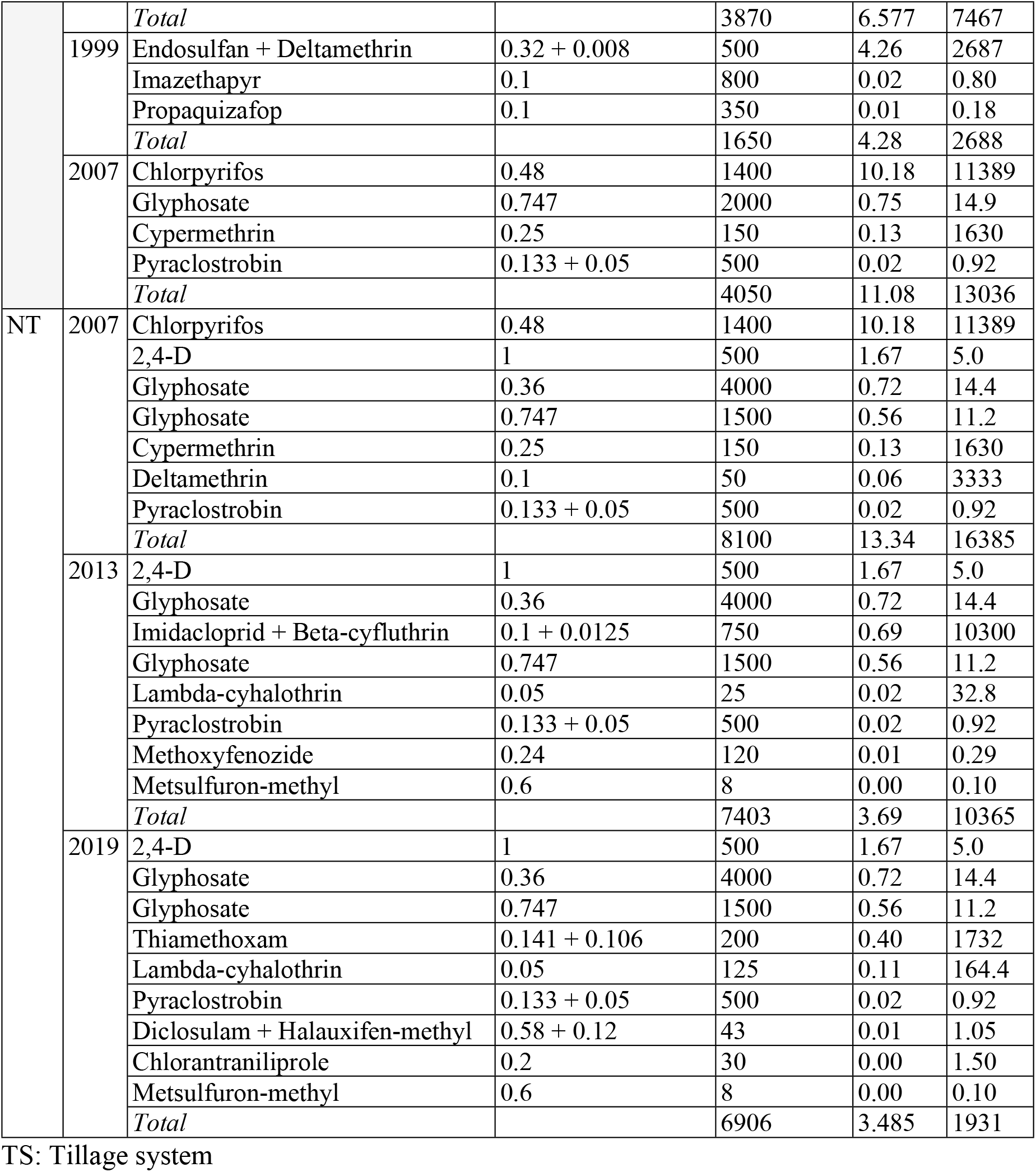
Toxic units for mammals (TUm) and insects (TUi) of pesticides used in soybean crop for the years 1987, 1993, 1999, 2007, 2013 and 2019. TS: Tillage system.

**Table 2.**
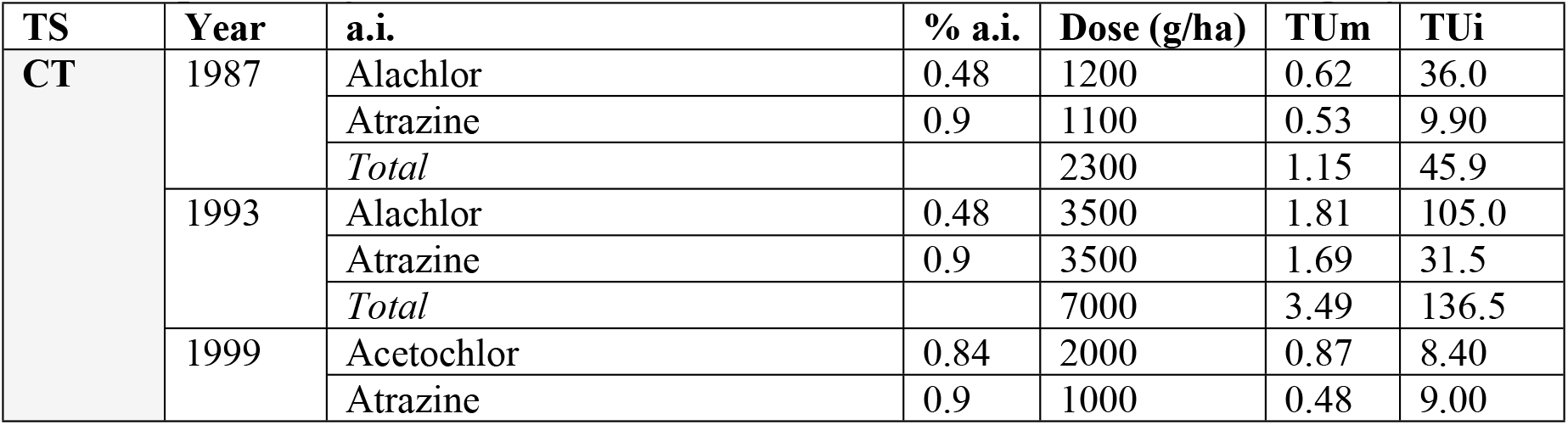
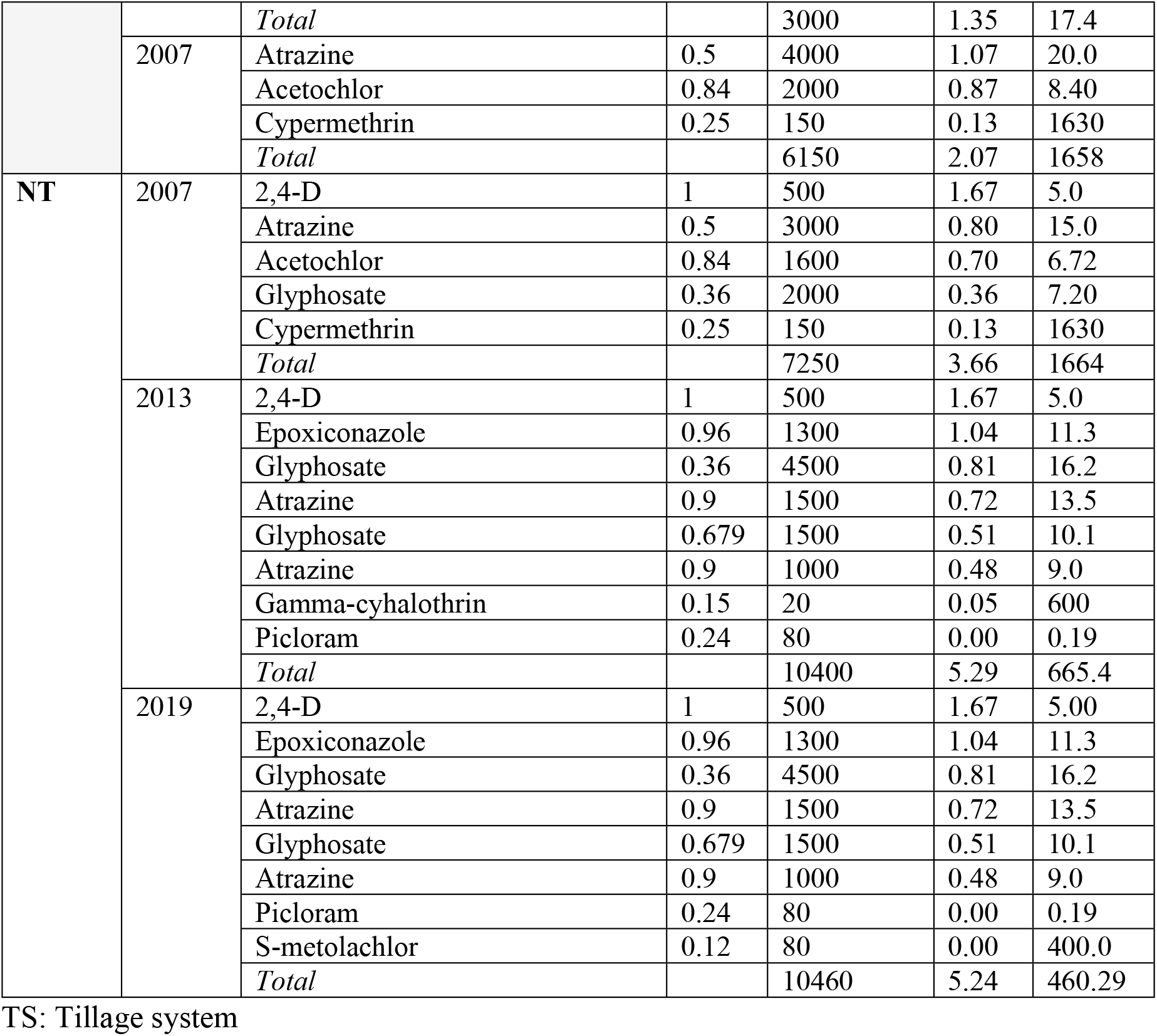
Toxic units for mammals (TU m) and insects (TU i) of pesticides used in maize crop for the years 1987, 1993, 1999, 2007, 2013 and 2019. TS: Tillage system.

Results from the studied period in the entire database of pesticide usage in soybean showed that in the first 20 years (1987-2007) the amount of pesticides applied ranges between 3000 and 4000 g/ha (Table 1). However, as a result of the shift to no-tillage system the applied dose doubled up to 8100 g/ha and has remained almost unchanged until 2019. The same pattern of increase in the amount of pesticide use was observed in maize crops, ranging from 2300 g/ha in 1987 up to 10460 g/ha by 2019 (Table 2). Regarding the ecotoxicity hazard of the pesticides used, the soybean showed no important changes in the total toxic units for insects (TUi) until the end of the period studied when final values were one order of magnitude lower than the rest of the database (Table 1). The mammal ecotoxicity assessment, measured in toxic units for mammals (TUm), showed the highest toxicity values at the beginning of the time series, mainly due to the used of an extremely toxic compound (Parathion-methyl) that was banned by 1990s. In the following years, the total number of TUm remained in the range of 5-13 TUm, until Chlorpyriphos and Endosulfan were replaced by less toxic insecticides (Table 1). In maize, the main change in insect ecotoxicity (TUi) was observed because of the adoption of insecticides in the analyzed pesticide profiles (Table 2). However, pesticide usage data showed a breakpoint toward lower doses and less toxic compounds in maize which resulted in fewer toxic units for insects by the end of the time series. Finally, mammal toxicity of pesticides usage in maize crop showed a constant and positive increase from 1.15 Tum, for the pesticide profile in 1987, up to 5.24 TUm in 2019 (Table 2).

## Discussion

Agricultural intensification relies on ecosystem assessment in order to move towards sustainable farming systems. However, this assessment should include both present and near future effects on key ecosystem processes in order to infer ecosystem time trends [38]. Regarding pesticide use, long-term monitoring represents a critical step for sustainability assessment as data on pesticide use remains scattered and not necessarily publicly available [39]. This is often the situation in developing countries where data on both pesticide monitoring and actual amount of pesticides use at national and regional are difficult to find [40]. In this paper, we cope with this problem by using standard pesticide regimes registered in one of the main cropping areas of Argentina as a proxy of the actual use of pesticides. This assumption may imply a significant caveat to be considered when analyzing our results, particularly when this regime is scaled-up using the sown area for assessing an overall pesticide hazard in maize and soybean crops. In this sense, the lack of direct information on the use of pesticides could incorporate some bias in the observed absolute values. However, both the data integrity on pesticide use regimes and the evolution of the sown area in both crops and under different tillage systems allows evaluating the observed trends and using them as reliable indicators of long-term change in the systems studied [41].

Environmental assessment should not only include the spatial and temporal dimensions, but it should also rely on meaningful metrics [42]. Concerning pesticide use, there are plenty of indicators based in commercialized volume, dose applied, exposure or several ecotoxicity values [43]. However, indicators based solely on the weight of pesticide applied can result in ambiguous or incorrect conclusions, because pesticides used in cropping systems involve a variety of toxicity profiles. This has led to a switch in current risk assessments from quantity-based [44] to toxicity-based indicators [45]. In this work, we used a toxicity-based indicator which follows a hazard quotient approach [12,46]. The selected indicator is free from most of the previous concerns about the environmental impact quotient (EIQ) developed previously [47]. Most of these concerns are related to numerical calculations with ordinal values, the undermining of important pesticide risk factors, the lack of supporting data for assigning some partial risk values and the strong correlation between field EIQ and pesticide use rate [12,15,16]. The acute toxicity on insect and mammals were integrated using fuzzy logic as a tool. The fuzzy logic approach has been previously used in pesticide assessment [18,48] and is very useful in order to develop a continuum process of ecosystem monitoring. The explicit nature of both membership functions and fuzzy if-then rules set up a conceptual assessment framework that could be easily improved in the future by the inclusion of new rules for weighting indicator scores in different situations [49]. This is a critical issue when modeling and building sustainability indicators as its characterization should be literal, and system-oriented [38]. Moreover, the scores derived from RIPEST involve the distance between observed values and some reference values rather than an absolute value, which rarely reveals whether the impact of a system is acceptable or not [50].

Our study is, as far as we know, the first long-term analysis of pesticide risk in cropping systems of Argentina. Previous long-term analysis showed that pesticide impact decreased in UK from 1992-2008, but also that this pattern is crop-dependent, with an initial risk decrease followed by a long stabilized period [11]. Data from herbicide use in USA from 1990-2015 showed that acute toxicity decreased for the six main crops; particularly both maize and soybean showed a decrease in acute toxicity per hectare [12]. Our results did not show a common time trend in ecotoxicity risk among crops. Results showed two temporal dynamics of pesticide impact, which were related to the analyzed crop. Soybean showed high temporal variability due to both technological changes associated to tillage system shift and a national ban of highly toxic compounds [51–53]. Pesticide regulations shows differences in Argentina in relation to EU countries mainly associated with the timing of prohibition and restriction (Iturburu *et al.*, 2019). However, several organophosphorus, all organochlorine pesticides (OCPs) have been restricted since 1991 and totally banned by 1998 [51,52,54]. By this time, the observed soybean time-trend showed a significant decrease both per area and total impact due to pesticide profiles as well as by the end of the studied period when impact reduction was mainly due to lower applied doses.

The Mann-Kendall statistical test only evaluates monotonic trends over the entire 32-year period. However, partial trends may be important, even where the overall trend is non-significant [12]. A partial positive trend in pesticide impact was observed both in soybean and maize crops during the period of no-tillage adoption. Pesticide risk increased in this period due to the double effect of the increase in the sown area and the shift towards systems with greater use of pesticides. When tillage is reduced, farmers become more reliant on other weed and pest control practices, and at least some of the widespread increase in pesticide use could be attributable to adoption of conservation tillage practices [55]. However, by the end of the time series (the period when no-tillage was fully adopted) the observed trend was different between crops. Although the sown area increased in both crops, total pesticide impact in these last years decreased in soybean because of a significant low pesticide use in the modal profiles. Otherwise, maize kept the toxicity profiles relatively constant during half of the time series, but the sown area increment resulted in a significant partial incremental pesticide impact trend. Soybean production showed a continuous improvement in its toxicity indicators, something opposite to what was observed in the corn crop. Maize has a continuous increased in ecotoxicity risk, boosted mainly by a higher dose trend. This result seems to be contrary to technological changes in maize, mainly represented by the incorporation of the genetic modification that confers resistance to insects (i.e. Bt-corn) as early as 1996 [56]. However, around the same period, herbicide resistance has been extensively documented in this productive area [57,58], which led to an increase in the use of herbicides [59]. This process was also enhanced by the relative increase in rotation of winter fallows without crop coverage due to the noticeable reduction of sown area with wheat and barley [60].

Environmental monitoring should encourage pesticide use changes towards more sustainable trajectories. However, the adoption of these changes depends on the way the observed trends are communicated and highlighting potential tradeoffs in pesticide assessment by using both impact and return metrics [61]. Ecoefficiency is a key indicator for showing an improved measure of sustainability because it links environmental impacts directly with some kind of economic performance, possibly leading towards sustainable development [62]. Time-trends of soybean showed a significant increase in ecoefficiency, particularly in the last years analyzed. Soybean dependence on herbicides has risen as a result of weed-related problems. However, RIPEST is sensible to acute toxicity defined mainly for highly toxic insecticides, and the modal pesticide soybean profiles showed a constant reduction not only in the number but also in the acute toxicity of insecticides used. Oppositely, the maize crop did not show improvements in ecoefficiency. As we previously mentioned, during this period some innovations have been adopted to increase yield and reduce the risk of crop loss [56]. These objectives were met as shown by the constant increase in total yield, remarkably during the las 10 years of the data analyzed. However, these improvements were not fully reflected in the ecoefficiency. This pattern is clearer when the ecoefficiency values are expressed in a common base, related to initial values. The constant value of relative ecoefficiency in maize crop is showing a possible path towards an unsustainable cropping system.

Finally, some areas for improvement in pesticide impact assessment has should not be overlooked. We reported data on environmental impact based on ecotoxicity. However, there are some issues to consider such as pesticide fate and transport, which is especially critical when assessing sensitive areas such watersheds and the urban-rural interface. However, data on long-term ecotoxicity trends, both on total impact and when using the ecoefficiency concept, should help to push for sustainable intensification by identifying negative trends and highlighting a potential tradeoff between crop productivity and environmental impact. The ultimate challenge is to reinforce high crop yield time-trends while simultaneously incentivizing the efficient use of pesticides to minimize this potential tradeoff between crop productivity and environmental impact.

## Acknowledgements

This material is based upon work supported by the University of Buenos Aires (UBA); the National Council for Scientific Research (CONICET); and the National Agency for Science Promotion (ANPCyT) of Argentina (PICT 2016-3216)

## Supporting Information

**S1 Table. Data for pesticide use**

**S2 Table. Data for sown area and yields**

